# Phenotypic analysis of various *Clostridioides difficile* ribotypes reveals consistency among core processes

**DOI:** 10.1101/2025.01.10.632434

**Authors:** Merilyn A. Beebe, Daniel Paredes-Sabja, Larry K. Kociolek, César Rodríguez, Joseph A. Sorg

## Abstract

*Clostridioides difficile* infections (CDI) cause almost 300,000 hospitalizations per year of which ∼15-30% are the result of recurring infections. The prevalence and persistence of CDI in hospital settings has resulted in an extensive collection of *C. difficile* clinical isolates and their classification, typically by ribotype. While much of the current literature focuses on one or two prominent ribotypes (*e.g*., RT027), recent years have seen several other ribotypes dominate the clinical landscape (*e.g.*, RT106 and RT078). Some ribotypes are associated with severe disease and / or increased recurrence rates, but why are certain ribotypes more prominent or harmful than others remains unknown. Because *C. difficile* has a large, open pan-genome, this observed relationship between ribotype and clinical outcome could be a result of the genetic diversity of *C. difficile*. Thus, we hypothesize that core biological processes of *C. difficile* are conserved across ribotypes / clades. We tested this hypothesis by observing the growth kinetics, sporulation, germination, bile acid sensitivity, bile salt hydrolase activity, and surface motility of fifteen strains belonging to various ribotypes spanning each known *C. difficile* clade. In viewing these phenotypes across each strain, we see that core phenotypes (growth, germination, sporulation, and resistance to bile salt toxicity) are remarkably consistent across clades / ribotypes. This suggests that variations observed in the clinical setting may be due to unidentified factors in the accessory genome or due to unknown host-factors.

**Importance:** *C. difficile* infections impact thousands of individuals every year many of whom experience recurring infections. Clinical studies have reported an unexplained correlation between some clades / ribotypes of *C. difficile* and disease severity / recurrence. Here, we demonstrate that *C. difficile* strains across the major clades / ribotypes are consistent in their core phenotypes. This suggests that such phenotypes are not responsible for variations in disease severity / recurrence and are ideal targets for the development of therapeutics meant to treat *C. difficile* related infections.

## Introduction

*Clostridioides difficile* is a Gram-positive, anaerobic, endospore forming pathogen with two major life stages: the metabolically active vegetative cell, and the dormant spore (1, 2). The spore is the transmissible form and provides extreme resistance to antibiotics, environmental stresses, and disinfection techniques (2–4). When a patient experiencing antibiotic-induced dysbiosis ingests *C. difficile* spores, the spores traverse the gut and germinate to the vegetative form (5–7). In the absence of a healthy microbiome, vegetative cells more efficiently colonize the host and can produce toxins which induce the symptoms characteristic of *C. difficile* infection (CDI) (diarrhea, pseudomembranous colitis, etc.) (8). These vegetative cells will form new spores which can remain in the gut or pass into the environment contributing to the spread / recurrence of disease (6). Of approximately 300,000 CDI related hospitalizations in the US, about 25-30% of patients experienced disease recurrence; the chance of recurrence increases with each subsequent infection (9, 10).

In 2006, the first fully assembled *C. difficile* genome was published (11). Currently, there are greater than 19,000 *C. difficile* genomes deposited in the NCBI database and greater than 31,000 deposited in Enterobase (12, 13). Broadly, *C. difficile* genomes are ∼4 Mbp in size and contain ∼4k genes (11, 13–15). About 25-50% of these genes belong to the core genome (genes shared among sequenced strains) while the pan-genome (all genes both shared and unique found in sequenced strains) is generally agreed to consist of ∼6k genes with no currently defined limit (16–18). In addition, greater than 11% of any given *C. difficile* genome consists of mobile elements (transposons, prophage, a skin element, etc.) (11). Phylogenetic analyses group *C. difficile* strains into five main clades and three cryptic clades (19, 20). Most studies agree that Clades 1 and 2 are closely related while Clade 5 exhibits the most genetic distinction, with recent studies suggesting that Clade 5 is undergoing speciation (18, 21). The relatively small size of the core genome, open pan-genome, large number of mobile elements, and observed evolutionary distance all indicate that *C. difficile*, as a species, has substantial genetic variation between its members.

In the clinical setting, *C. difficile* strains are typically classified using various typing methods (ribotyping, restriction endonuclease analysis [REA], multilocus sequence typing [MLST], toxinotyping, serotyping, etc.) (22–27). Of these, ribotyping remains the most popular and clinically relevant. Historically, ribotype 027 (RT027) strains have been associated with the worst CDI outbreaks, and thus are generally the most well studied (14, 15, 28, 29). However, the prominent strains isolated from more recent outbreaks belong to less well-studied ribotypes (*e.g.*, RT106 or RT078) (30–38). Because *C. difficile* is so genetically diverse, ribotype-specific clinical outcomes may be attributable to processes encoded by the accessory genome and not to functions encoded in the core-genome. Thus, we hypothesize that processes central to *C. difficile* biology (*e.g.,* vegetative growth, sporulation, germination, and resistance to bile salt toxicity) are conserved across ribotype / clade. To test this hypothesis, we collected three strains from one ribotype belonging to each of the five classical clades. We then measured their growth in rich and minimal media, spore production, response to germinants, bile salt hydrolase activity, resistance to bile acid toxicity, and surface motility. Our analyses indicate that, while the strains show variations in their ability to process taurocholate-conjugated bile salts, they exhibit remarkably consistent core phenotypes with no strong patterns across ribotypes / clades. These results support the hypothesis that despite the evolutionary variation observed between strains of *C. difficile*, certain processes that are central to *C. difficile* biology remain consistent across ribotype / clade.

## Methods

### Bacterial strains and growth conditions

*C. difficile* R20291 was used as an experimental control for all assays performed in this study. *C. difficile* strains LC5624, LK3P-030, and LK3P-081 were obtained from pediatric patients receiving care at the Ann & Rober H. Lurie Children’s Hospital of Chicago, Chicago, IL, USA. *C. difficile* strains M68 and M120 were obtained from the collection of Dr. Trevor Lawley (Sanger Institute, Hinxton, UK). The remaining strains were collected by Dr. Daniel Parades-Sabja (Texas A&M University, College Station, TX, USA) and César Rodríguez (Facultad de Micriobiología & Centro de Inventigación en Enfermedades Tropicales, Universidad de Costa Rica, San José, Costa Rica) from various outbreaks in Central / South America.

Strains were routinely grown anaerobically (Coy Laboratories, model B, 4% H_2_, 5% CO_2_, 85% N_2_) at 37 °C. Each strain was grown on / in either brain heart infusion medium supplemented with yeast extract and 0.1% (w / v) L-cysteine (BHIS) or in *C.* difficile minimal medium (CDMM; 1% (w / v) casamino acids, 2 mM L-tryptophan, 4 mM L-cysteine, 35.2 mM Na_2_HPO_4_ * 7H_2_O, 60 mM NaHCO_3_, 7 mM KH_2_PO_4_, 20 mM NaCl, 45 mM glucose (or indicated carbohydrate), 400 μM (NH_4_)_2_SO_4_, 177 μM CaCl_2_ * 2H_2_O, 96 μM MgCl_2_ * 6H_2_O, 50.6 μM MnCl_2_ * 4H_2_O, 4.2 μM, CoCl_2_ * 6H_2_O, 34 μM FeSO_4_ * 7H_2_O, 4 μM D-biotin, 2 μM calcium-D-pantothenate, 6 μM pyridoxine). Where indicated, the pH of BHIS was adjusted using acetic acid / sodium hydroxide or supplemented with 0.1% taurocholate (TA; GoldBio S-121-100) in solid medium and 1 mM TA, taurodeoxycholic acid (TDCA; Millipore Sigma 580221-50GM), hyodeoxycholic acid (HDCA; MP Biomedicals 157514), cholic acid (CA; Sigma Aldrich C1129-100G), chenodeoxycholic acid (CDCA; ACROS Organics RK-88245-39), or deoxycholic acid (DCA; Sigma Aldrich D2510-100G) for liquid cultures. Except for TA, which is soluble in water, bile acids stocks were made in dimethylsulfoxide (DMSO).

### Whole Genome Sequencing

Genomic DNA was extracted for all strains except *C. difficile* R20291 (GenBank accession number NZ_CP029423.1), *C. difficile* LC5624 (GenBank accession number CP022524.1), *C. difficile* M68 (GenBank accession number NC_017175.1), and *C. difficile* M120 (GenBank accession number NC_017174.1) for which genomes are already published. Briefly, each strain was grown for 18 hours in BHIS medium. The cells were pelleted, lysed, and subjected to phenol-chloroform extraction. This genomic DNA was sent to SeqCoast Genomics (Portsmouth, NH, USA) for long-read Oxford Nanopore sequencing. The long-read data was assembled using the Flye plugin for *de novo* assembly in Geneious Prime (version 2024.0.5) (39, 40). To improve genome quality, Illumina sequencing reads for each of the strains were then mapped to the assembled genomes using BBMapper (41). The genome alignments were performed using the MAUVE plugin (42). Phylogenies were generated using the Geneious Tree Builder with a Tamura-Nei genetic distance model and the neighbor-joining method for three individual locally colinear blocks (LCBs) produced by the MAUVE alignment.

### Protein alignment

The genes encoding each of the indicated proteins were extracted from the assembled and MAUVE aligned genomes. Their sequences were translated and aligned using ClustalOmega in GeneiousPrime (43). The amino acids were shaded according to similarity (Pam140 Score Matrix) with white indicating 100% amino acid identity across all observed strains and black indicating <60% identity.

### Growth curves and doubling times

For growth curves in BHIS medium, strains were grown in pre-reduced BHIS liquid medium for 12 hours. Cultures were back diluted to an OD_600_ = 0.05 and allowed to grow to an OD_600_ = 0.5. This log-phase culture was then used to inoculate the experimental culture to an OD_600_ of 0.05 at T_0_. OD_600_ measurements were taken every 30 minutes for 8 hours using a Biowave Cell Density Meter CO8000. When the OD_600_ reached ≥0.3, the OD measurements across 2 hours of growth were used to calculate doubling time.

For growth curves in CDMM, strains were grown in pre-reduced CDMM (made with a final concentration of either 45 mM glucose / fructose, 53 mM xylose, or 50 mM trehalose, where indicated) for 24 hours. Cultures were back diluted to an OD_600_ = 0.05 and grown for 12 hours to an OD_600_ ≈ 0.5 – 0.7. The culture was then used to inoculate 3 mL of the indicated medium to a starting OD_600_ of 0.05 in a 12-well plate. The plate was then placed in a Stratus Microplate Reader (Cerillo, Charlottesville, VA, USA) where OD_600_ measurements were collected every 3 minutes for 22 hours. Each time point represents the average reading from three different sensors measuring a single well. Due to random blockage of sensors during the assay, the data was filtered using the z-score method. Doubling times were calculated using the most linear portion of the OD curves and the formula t_2_ = ln(2)/(k) where k is the growth rate determined using the formula k = LN(OD_t2_/OD_t1_)/(t2-t1).

### Spore purification

Spores were purified as previously described (44–51). Briefly, the indicated strains were plated onto 10 pre-reduced BHIS plates and incubated for 5 days. The growth from each plate was scraped into 1 mL 18 MΩ dH_2_O and stored at 4 °C. After a minimum period of 24 hours, cells were resuspended in 1 mL 18 MΩ dH_2_O and centrifuged for 1 minute at 14,000 x g. The supernatant was removed, and subsequent washes with 18 MΩ dH_2_O performed, separating the spores from vegetative cells / cell debris in distinct layers which were gradually removed and cell pellets combined with each wash. Final contaminants were removed by placing the cells on 9 mL of 50% sucrose and centrifuging 20 minutes at 4,000 x g and 4 °C. The supernatant was discarded, and the pellet (containing the purified spores) was washed thrice to remove the remaining sucrose. The final pellet was resuspended in 1 mL 18 MΩ dH_2_O and stored at 4°C.

### Sporulation

The sporulation assay was performed as previously described, with some slight modifications (3, 52). A 16 hour culture of the indicated strain was back diluted to an OD_600_ = 0.05 and allowed to grow to an OD_600_ = 0.5. One hundred microliters of the log-phase culture was plated on pre-reduced BHIS agar medium and incubated for 48 hr. One quarter of the plate was scraped into 1 mL of 1x PBS (pH 7.4) and 500 μL of this suspension was treated with 100% ethanol for 20 minutes. The treated cells were serially diluted in 1x PBS with 0.1% TA and plated on pre-reduced BHIS TA plates. The plates were incubated for 48 hours prior to CFU enumeration.

### Germination assay

Germination was assessed using an OD-based assay as described previously (45, 46). Briefly, samples of purified spores were adjusted to an OD_600_ of 0.5. The spores were heat treated at 65 °C for 30 minutes. For each of the tested germinant concentrations, 5 μL of the spore sample was suspended in 95 μL of the appropriate germination buffer. For all samples the buffer contained 0.5 M HEPES, 50 mM NaCl, pH 7.2. For co-germinant (glycine) efficiency, the buffer contained 10 mM TA and various amounts of glycine (final concentrations of 0 mM, 0.1 mM, 0.2 mM, 0.4 mM, 0.6 mM, 0.8 mM, 1 mM, 2 mM, or 5 mM). For bile-acid germinant (TA) efficiency, the germination buffer contained 100 mM glycine and 10% (v / v) DMSO (to control for the DMSO used to dissolve CDCA), and various amounts of TA (final concentrations of 0 mM, 0.1 mM, 0.2 mM, 0.5 mM, 1 mM, 2 mM, 5 mM, or 10 mM). For CDCA, the germination buffer was supplemented with 100 mM glycine and 1 mM CDCA and TA to a final concentration of 1 mM, 2 mM, 5 mM, 10 mM, 15 mM, 20 mM, 30 mM, or 50 mM. The OD_600_ of each sample was recorded every 30 seconds over the course of 1 hour. The plate was shaken vigorously 5 seconds before each measurement.

A germination curve was generated for each strain / germinant combination by plotting the OD_600_ at a given time (T_x_) divided by the OD_600_ at time zero (T_0_) vs. time (the controls for all strains / germinants and an example for R20291 with varying concentration of glycine are shown in Figure S1) (45, 46, 53–56). Germinant sensitivity was calculated using the maximum slope for each germination curve. The slope was plotted against (co)-germinant concentration to generate a Michaelis-Menten graph. A Lineweaver-Burke plot was generated, and from this, the Ki/EC_50_ was calculated. Here, EC_50_ is defined as the concentration of germinant which produces half the maximum germination rate. The efficiency of the competitive inhibitor was calculated as previously described using the following equation K_i_ = [inhibitor] / ((K_CDCA_ / K_TA_)-1) (53–56).

### Bile salt sensitivity

Each strain was grown for 16 hours, back diluted to an OD_600_ = 0.05, and allowed to grow to an OD_600_ = 0.5. Fifty microliters of these cultures were added to pre-reduced BHIS of the indicated pH and bile acid concentration (obtained through a series of 1:1 dilutions to a final volume of 500 μL). The samples were incubated for ∼18 hours and MICs were assessed by the presence / absence of growth.

### Bile salt hydrolase activity

Bile salt hydrolase activity assays were performed as described previously (48). Briefly, strains were grown in BHIS liquid medium for 16 hours. 10^8^ CFU from these cultures were transferred into 5 mL BHIS supplemented with 1 mM TA or TDCA and incubated for 24 hours after which the cultures were centrifuged for 10 min at 4,000 x g. The pellet was suspended in 100% methanol. One millimolar HDCA (an internal standard) was added to the supernatant before being lyophilized and suspended in methanol. The suspended pellet and dried supernatant were combined. Each strain was run alongside three *C. difficile* R20291 controls: a negative control without TA, a control with 1 mM each TA / TDCA, CA / DCA, and HDCA all added to the spent supernatant, and a positive control with 1 mM TA added before the 24 hour incubation (Figure S2).

The bile salts found in each sample were separated by reverse-phase high performance liquid chromatography (HPLC) using a Shimadzu Prominence system (Shimadzu, Kyoto, Japan) (48, 57–59). For each strain, 30 μL of the sample was separated on a Synchronis C18 column (4.6 by 250 mm, 5 μm particle size, ThermoFisher, Waltham, MA, USA) using a methanol-based mobile phase [53% methanol, 24% acetonitrile and 30 mM ammonium acetate (pH 5.6)]. A Sedere Sedex model 80 low temperature-evaporative light scattering detector (LT-ELSD) using 50 psi Zero Grade air at 94 °C detected the bile salt peaks. Percent deconjugation was calculated using the area under the peak for CA / DCA divided by the sum of the areas under the peaks for TA / TDCA and CA / DCA.

### Surface Motility

Surface motility assays were performed as previously described (60). Briefly, strains were grown in BHIS for 16 hours, diluted 1:50 in fresh BHIS medium, and grown until the OD_600_ reached ∼0.5. From these cultures, 10 μL was spotted onto pre-reduced BHIS agar medium and incubated for 5 days. Plates were then imaged using a Bio-Rad GelDoc XR+ (Bio-Rad Laboratories, Hercules, CA, USA).

### Statistical analysis

All data represent the average from three independent biological replicates with the error bars indicating the standard error of the mean. For each, statistical significance was determined using the one-way ANOVA analysis function from GraphPad Prism (version 9.0.2 for Windows, GraphPad Software, San Diego, California USA). When results were compared to *C. difficile* R20291, a Šidák’s multiple comparisons test was used, while a Tukey’s multiple comparisons test was used when comparing between ribotypes / clades. Asterisks indicate p-values with, * = < 0.5, ** = < 0.02, *** = < 0.01, and **** = < 0.0001.

## Results

### Strain collection and genomic analysis

To study phenotypic variation among the *C. difficile* clades, we collected 15 different clinical isolates of *C. difficile* (Table 1). Each of the five main clades are represented in this study by three distinct strains belonging to a single ribotype within the clade. The indicated ribotype was selected based on its clinical relevance (30, 37, 61–64). Apart from those strains whose genomes have already been published (*C. difficile* R20291, *C. difficile* LC5624, *C. difficile* M68, and *C. difficile* M120), we assembled the genomes for each of the strains and deposited them in the NCBI database. Phylogenetic analysis clusters the strains into the 5 classical clades, consistent with previously published data (Figure 1, S3) (18, 21, 65).

**Figure 1:**
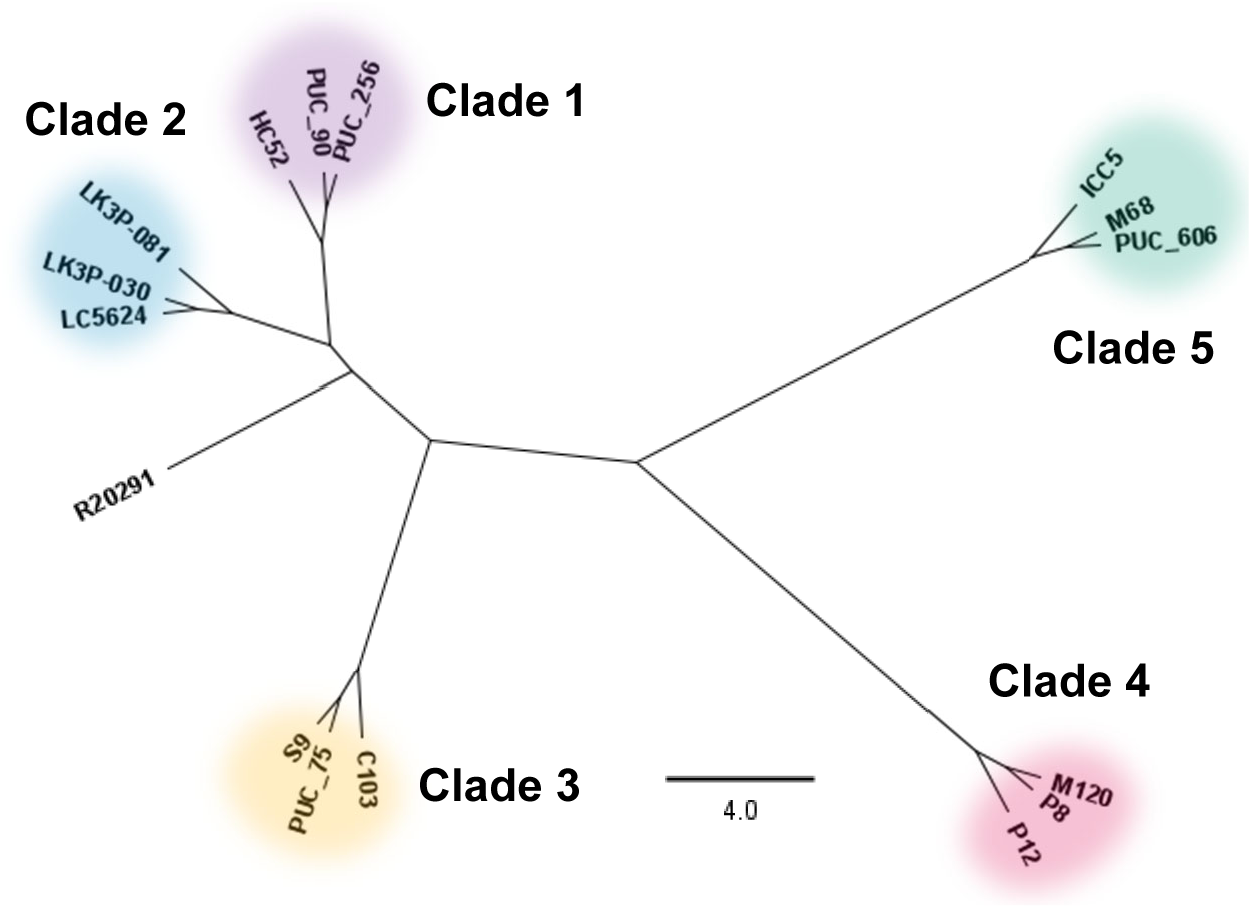
Phylogeny of strains used in this study. The neighbor joining phylogeny generated for the strains in this study created from LCB 145, a 1,418,215 bp segment of the MAUVE alignment representing approximately 25% of any given genome in the study. The phylogeny was created using the Geneious Tree Builder application in the Geneious Prime software using the Tamura-Nei genetic distance model. Strains are grouped by their respective ribotypes / clades with the scale bar representing the number of substitutions per 1000 bp.

**Table 1:** Strains used in this study.

| Strain | Clade | Ribotype |
| --- | --- | --- |
| R20291 | 2 | 027 |
| PUC_256 | 1 | 014-020 |
| HC52 | 1 | 014-020 |
| PUC_90 | 1 | 014-020 |
| LC5624 | 2 | 106 |
| LK3P-030 | 2 | 106 |
| LK3P-081 | 2 | 106 |
| PUC_75 | 3 | 023 |
| S9 | 3 | 023 |
| C103 | 3 | 023 |
| M68 | 4 | 017 |
| PUC_606 | 4 | 017 |
| ICC5 | 4 | 017 |
| M120 | 5 | 078 |
| P8_S146 | 5 | 078 |
| P12_S145 | 5 | 078 |

### *C. difficile* growth is consistent in rich medium

In the laboratory setting, *C. difficile* is often cultured in a rich medium (BHIS). Because this was the medium in which all our assays would be performed, we first sought to determine if there were any inherent growth differences in this medium. Growth curves in BHIS medium were obtained for each strain and indicate little variation in overall growth kinetics. We observed minimal differences between each strain in a given clade (Figure 2A – E). All strains reached stationary phase within 4 hours of growth and a maximum OD_600_ of 2.0 – 3.0. Moreover, there were no significant differences in growth when the data were grouped by clade (Figure 2F). For a more objective comparison, we determined the generation times for each strain. The generation times for individual strains, including the *C. difficile* R20291 control, were calculated at 40 – 60 minutes (Figure 2G). The average generation time for each Clade was ∼50 minutes (Figure 2H). Taken together, this indicates that none of the tested strains exhibited a fitness advantage / disadvantage in BHIS medium.

**Figure 2:**
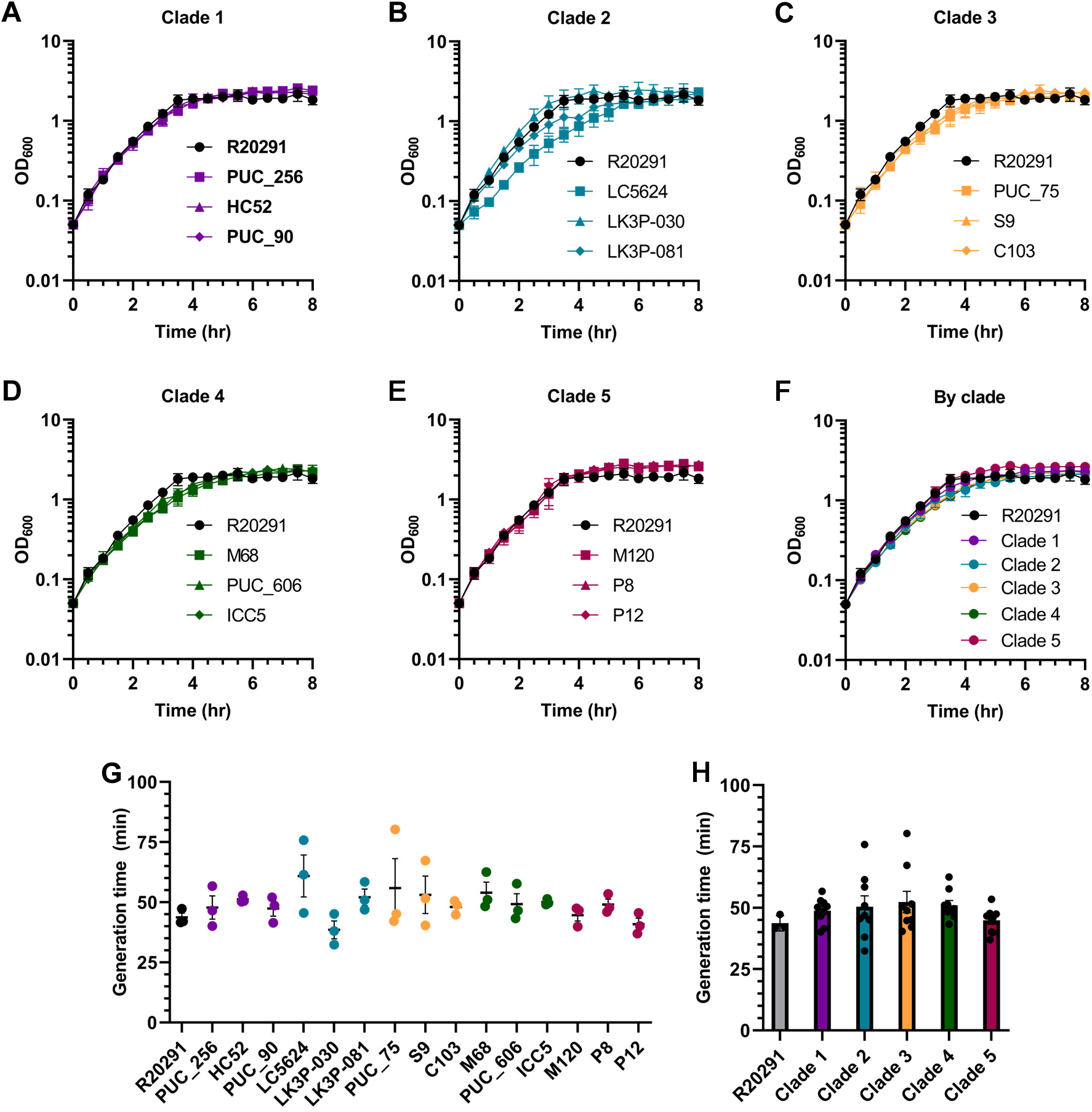
Growth of strains in rich medium. OD_600_ measurements for Clade 1 **(A)**, Clade 2 **(B)**, Clade 3 **(C)**, Clade 4 **(D)**, and Clade 5 **(E)** strains were taken every 30 minutes over the course of 8 hours. This same data is shown in **(F)** grouped by ribotype / clade. Data from the most linear portion of the growth curve was used to calculate doubling time presented by strain **(G)** and by ribotype / clade **(H)**. Data points represent the average from independent biological triplicates with error bars representing the standard error of the mean. For **(G)**, Šidák’s multiple comparisons test was used, while a Tukey’s multiple comparisons test was used for **(H)**. No statistically significant differences between strains were found.

### Clade 4 strains are limited in their ability to use various carbohydrates

Given that BHIS is a rich medium, the lack of variation in growth between strains was not entirely surprising. To determine if the strains responded differently in minimal medium, we cultured them in standard CDMM (containing glucose). Under these conditions, all strains exhibited less growth compared to growth in BHIS medium as indicated by noticeably increased generation times (Figure 3A). While the *C. difficile* R20291 control strain had a generation time of 100 minutes, *C. difficile* strains LK3P-081, C103, M68, PUC_606, ICC5, M120, and P12 had slower generation times. The Clade 4 strain, *C. difficile* M68, had the slowest generation time at ∼400 minutes (Figure 3B). When this data is grouped by clade, the Clade 1 strains had a generation time of ∼90 minutes while the Clade 4 strains had a generation time of ∼350 minutes (Figure 3C). Taken together, the data indicate that Clade 4 strains have a fitness disadvantage in standard CDMM relative to the other strains.

**Figure 3:**
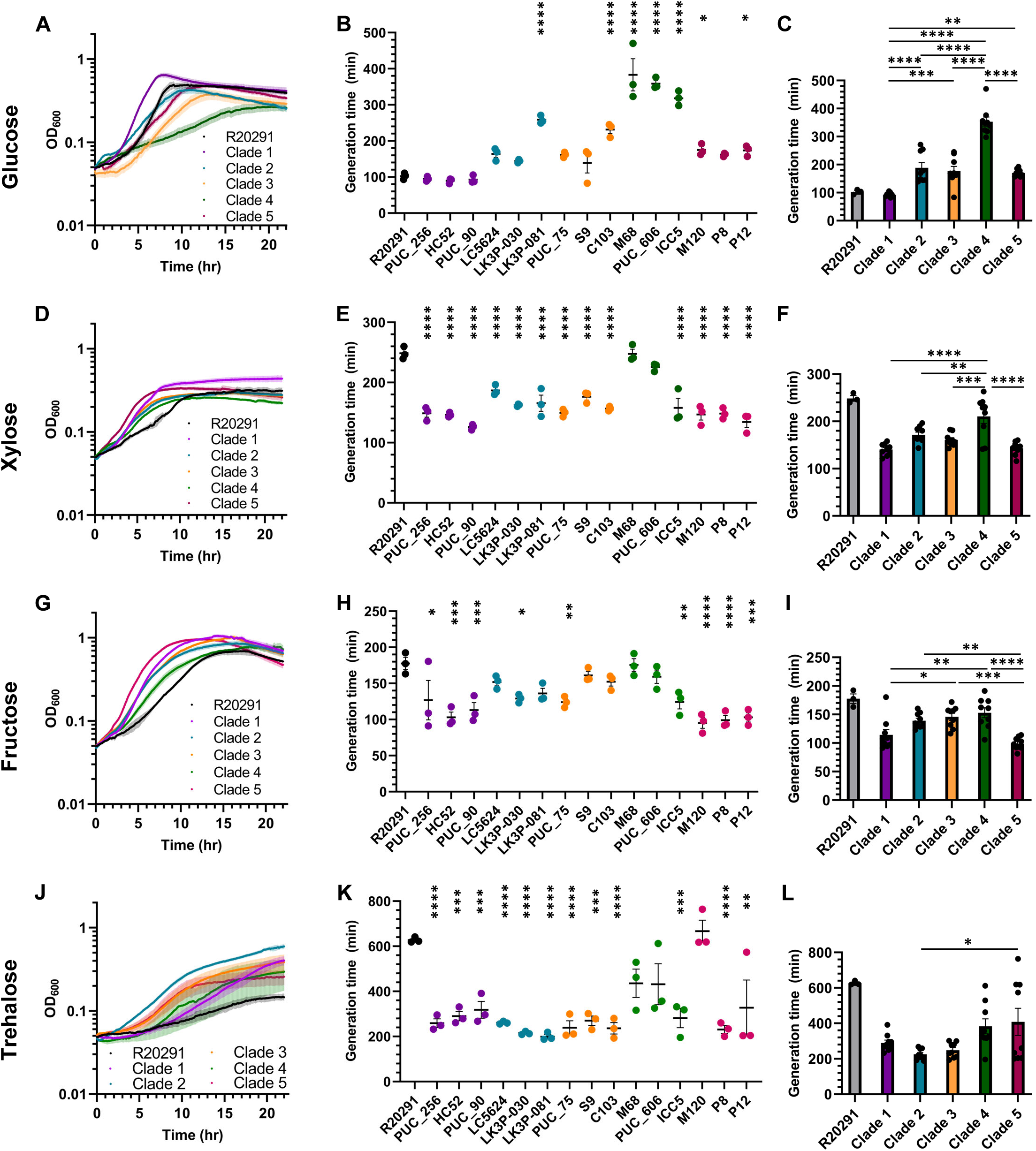
Growth of strains in minimal medium. Strains were grown in CDMM supplemented with either glucose **(A)**, xylose **(D)**, fructose **(G)**, or trehalose **(J)**. OD_600_ measurements for each strain were taken every 3 minutes for 22 hours. Data from the most linear portion of the growth curve was used to calculate generation times for growth in CDMM supplemented with glucose **(B,C)**, xylose **(E,F)**, fructose **(H,I)**, or trehalose **(K,L)**. Data points represent the average from independent biological triplicates with error bars representing the standard error of the mean. For **(B, E, H, and K)**, Šidák’s multiple comparisons test was used, while a Tukey’s multiple comparisons test was used for **(C,F,I, and L)**. Asterisks indicate p-values of * = < 0.5, ** = < 0.02, *** = < 0.01, and **** = < 0.0001.

Because *C*. *difficile* can incorporate different carbohydrates into its metabolic pathways, we sought to determine if any of the strains favored one carbohydrate source over another. Previous research has identified xylose as an important carbon source for many bacteria (66, 67). Xylose forms a five-carbon, six-member ring like glucose, but lacks a 6’ carbon and associated hydroxyl group. When testing growth in CDMM supplemented with xylose (CDMM-xyl) we observed slight variations between strains. All strains reached stationary phase between 6 – 10 hours (Figure 3D).. Contrary to what was observed for growth in standard CDMM, generation times for all strains except the *C. difficile* R20291 control (246 minutes), *C. difficile* M68 (240 minutes), and

*C. difficile* PUC_606 (220 minutes) were below 200 minutes (Figure 3E). Additionally, while most of the strains grew better than the control in CDMM-xyl, Clade 4 strains, once again, appeared to have a small fitness disadvantage in this medium in comparison to the other clades with an average doubling time of ∼210 minutes (Figure 3F).

Fructose, another common carbohydrate that is important for *C. difficile* metabolism, is the primary carbohydrate found in medium which allows for selection / isolation of *C. difficile* in clinical environments (TCCFA) (68, 69). Like glucose, it contains six carbons but forms a five-member ring. When grown in CDMM supplemented with fructose (CDMM-fru), the strains reached stationary phase between 8 – 12 hours (Figure 3G). *C. difficile* strains LK3P-030, PUC_75, ICC5 and all Clade 1 and 5 strains exhibited generation times of 100 – 130 minutes. These times were lower than the *C. difficile* R20291 control strain which had a 177-minute generation time (Figure 3H). *C. difficile* strains LC5624, LK3P-081, S9, C103, M68, and PUC_606 all grew similarly compared to *C. difficile* R20291 with generation times between 130 and 175 minutes. When grouped by clade, strains from Clades 2, 3, and 4 showed a ∼50-minute slower generation time relative to the remaining strains (Figure 3I).

Finally, we tested growth of the strains in CDMM supplemented with trehalose (CDMM-tre). Trehalose is composed of two α,α-1,1-linked glucose molecules and has, arguably, been associated with an increase in CDI (21, 70, 71). Growth in CDMM-tre was variable with Clade 2 strains reaching stationary phase around 10 hours of growth and Clade 1 strains reaching stationary phase around 20 hours of growth (Figure 3J). All strains, except for *C. difficile* M120 (660 minutes), had a lower generation time than the *C. difficile* R20291 control strain (620 minutes) (Figure 3K). Additionally, generation times for *C. difficile* strains PUC_256, HC52, PUC_90, LC5624, LK3P-030, LK3P-081, PUC_75, S9, C103, ICC5, P8, and P12 were similar to the generation times observed in CDMM-xyl at ∼200 – 300 minutes. Overall, Clade 4 strains had consistently slower growth in CDMM-tre relative to the other clades (Figure 3L).

### Variations in carbohydrate metabolism protein sequences are not consistent with phenotypic differences

The catabolite control protein, CcpA, is encoded by all strains in this study (Figure S4). Because there was little variation in the CcpA protein sequence, we hypothesize that there should be no major change in the global regulation of carbohydrate metabolism. In all tested strains, the xylose utilization operon was also present. Some variants appeared in the XylA (isomerase), XylB (kinase), and XylR (transcriptional regulator) protein sequences (Figure S5 – 7). However, none of these variations were consistent between the Clade 1, 2, 3, and 5 strains which had faster generation times compared to the *C. difficile* R20291 control strain (Figure 3D – F).

Previous reports found heterogeneity in the genes responsible for processing trehalose and corresponding differences in the ability of strains from different ribotypes to grow in media in which trehalose was the sole carbon source (21, 71). We observed the same for the strains tested herein, specifically when comparing the phosphotrehalase (TreA) protein sequence (Figure S8). *C. difficile* strains PUC_75 and S9 had nonsense mutations in TreA, and *C. difficile* strains C103, M120, P8, and P12 were missing TreA at this locus entirely. The trehalose operon repressor (TreR) also showed some variations and was missing entirely at the canonical locus in the Clade 3 and 5 strains (Figure S9). Both the *treR* and *treA* genes were found in an operon elsewhere in the genomes for the Clade 5 strains and these copies are included in the alignments (Figure S8 – 9). Clade 3 strain, *C. difficile* C103, contained an intact copy of *treA* at an additional locus, but did not possess a corresponding copy of *treR*, suggesting that *treA* may be regulated differently in this strain. The remaining Clade 3 strains (*C. difficile* PUC_75 and S9) do not possess an intact copy of *treA* suggesting the existence of another, non-canonical, trehalose metabolism pathway in these strains.

### Spore production is consistent between strains

Given that *C. difficile* is transmitted by spores, changes in sporulation levels could explain why some strains are more prevalent in the clinical setting than others; an increase in spore number may lead to increased disease spread / recurrence. We measured sporulation in each of the strains over a 48-hour period (Figure 4A – E) and observed a small increase in spore number for the Clade 1 strain *C. difficile* HC52 (∼10^9^ spores), and a decrease for the Clade 2 strain *C. difficile* LK3P-030 (∼10^7^ spores), relative to the *C. difficile* R20291 control (∼10^8^ spores). When this data was grouped by clade, the Clade 1 strains produced ∼10-fold more spores on average than the Clade 4 strains (Figure 4F).

**Figure 4:**
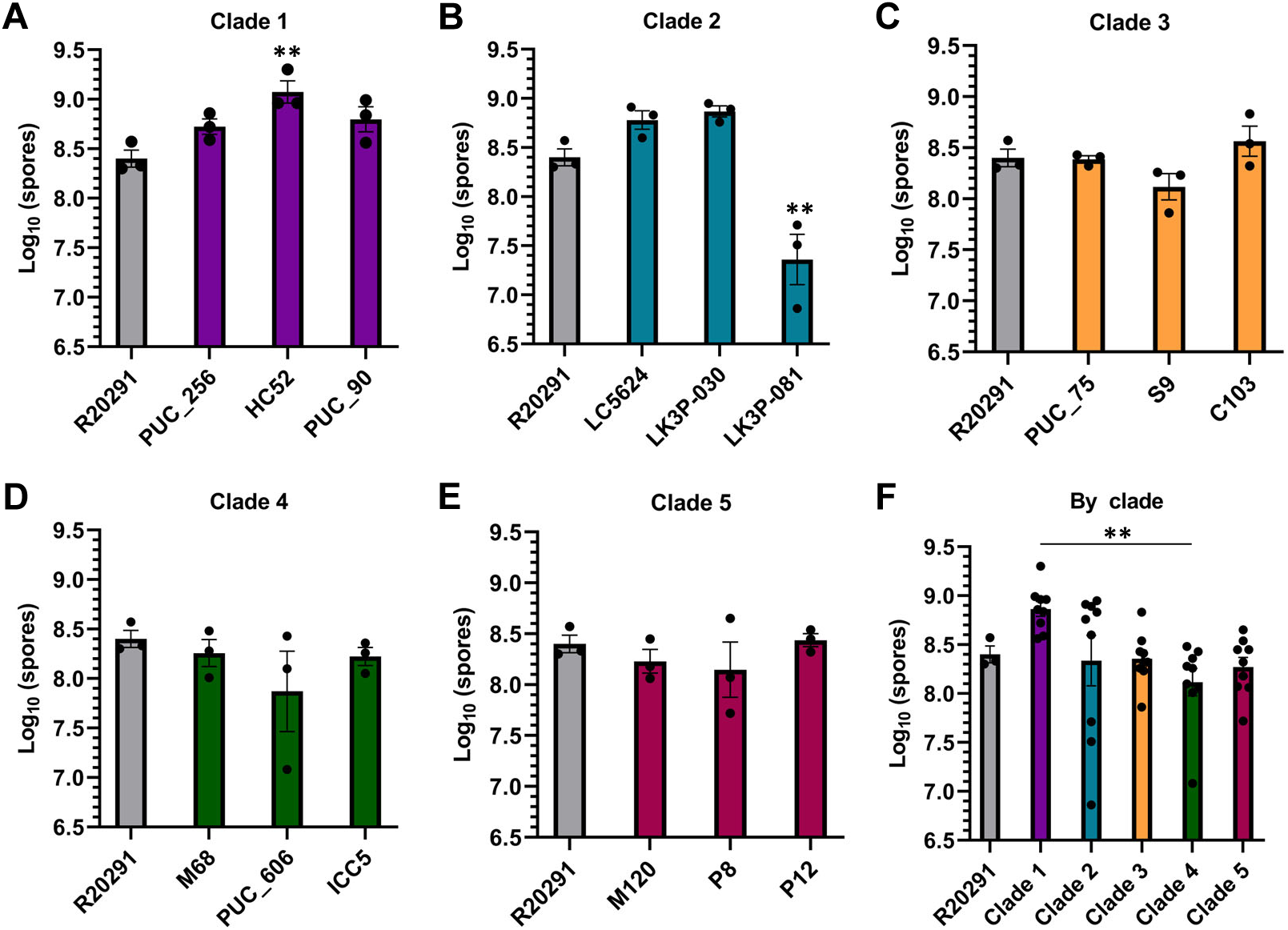
Spore production by strains over 48 hours. The number of spores produced on BHIS over 48 hours for Clade 1 **(A)**, Clade 2 **(B)**, Clade 3 **(C)**, Clade 4 **(D)**, and Clade 5 **(E)** reported on a log_10_ scale. This same data is grouped by clade in **(F)**. Data points represent the average from independent biological triplicates with error bars representing the standard error of the mean. For **(A – E)**, Šidák’s multiple comparisons test was used, while a Tukey’s multiple comparisons test was used for **(F)**. Asterisks indicate a p-value of ** = < 0.02.

### *C. difficile* clinical isolates are more sensitive to germinants than the *C. difficile* R20291 lab strain

Bile salts are cholesterol derivatives found in mammalian digestive systems and contribute to the absorption of nutrients (72). These compounds are produced in the liver and circulate through the small intestine before being recycled back to the liver. A small portion of these bile acids escape enterohepatic recirculation and enter the colon (73, 74). Germination by *C. difficile* spores occurs in response to two signals, a bile acid germinant and an amino acid co-germinant (75). Prior work has shown that TA and glycine are the most efficient germinants *in vitro* and consequently, they are used frequently in germination analyses (45). Because germination is required for successful outgrowth of vegetative cells from the dormant spore form, we sought to determine how the strains responded to germinants. Previous work from our lab and others has demonstrated that germination efficiency could be quantified as an EC_50_ value (53–56).

All strains except for *C. difficile* PUC_90 (Clade 1), had a lower EC_50_ value for glycine compared to the *C. difficile* R20291 control, which had an EC_50,glycine_ of 0.25 mM (Fig 5A). Clade 1 strains showed the most variation in EC_50,glycine_ with values ranging from 0.06 mM (*C. difficile* HC52) to 0.30 mM (*C. difficile* PUC_90). Clade 4 strains were the most consistent with EC_50,glycine_ values ranging from 0.05 mM – 0.08 mM. The average EC_50,glycine_ value for each clade fell between 0.07 mM and 0.18 mM indicating that glycine sensitivity is consistent between the strains (Figure 5B). The EC_50,TA_ value for all strains ranged from 0.10 mM (*C. difficile* P8) to 0.60 mM (*C. difficile* HC52), all of which were lower than the *C. difficile* R20291 control (2.0 mM) (Figure 5C). The largest variation was observed in the Clade 1 strains with EC_50,TA_ values ranging from 0.20 mM (*C. difficile* PUC_90) to 0.80 mM (*C. difficile* HC52). Clade 5 strains had the least variation with the EC_50,TA_ values ranging from 0.10 mM (*C. difficile* P8) to 0.15 mM (*C. difficile* P12). At the clade level, EC_50,TA_ remained consistent with only slight variations between 0.10 mM for Clades 3 and 5 and 0.60 mM for Clade 4 (Figure 5D).

**Figure 5:**
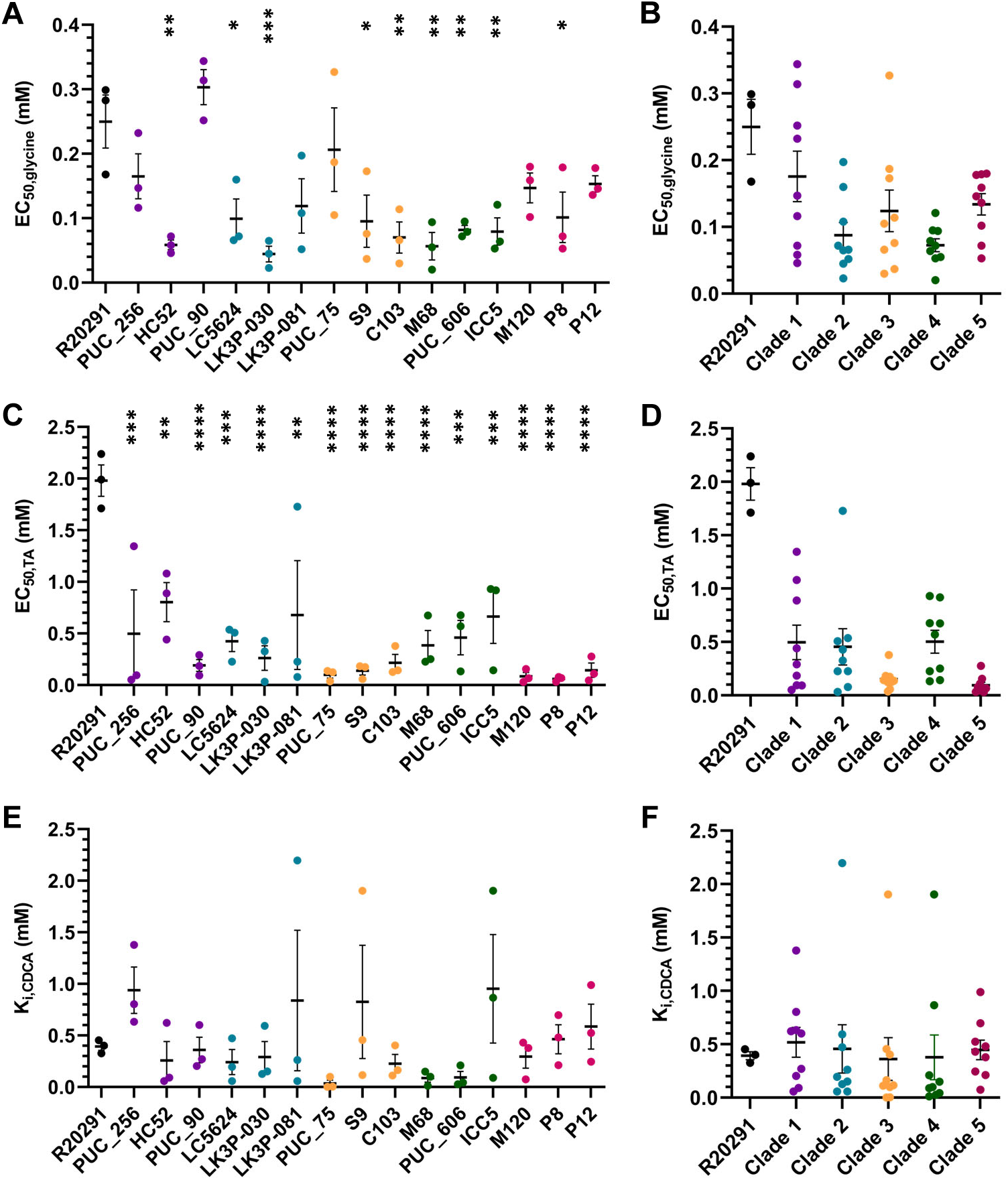
Strain sensitivity to germinants. Germination assays for each strain were performed in the presence of various concentrations of glycine **(A,B)**, TA **(C,D)**, or TA+CDCA **(E,F)**. Germinant sensitivity was calculated using the maximum slope for each condition plotted against (co)-germinant concentration. The data fitted to a linear relationship by taking the inverse of the slope vs. concentration plot and from this Ki/EC_50_ was calculated with EC_50_ equaling the concentration of germinant which produces the half maximum germination rate. The efficiency of the competitive inhibitor was calculated using the following equation K_i_ = [inhibitor] / ((K_CDCA_ / K_TA_)-1) (55, 56). Data points represent the average from independent biological triplicates with error bars representing the standard error of the mean. For **(A, C, and E)**, Šidák’s multiple comparisons test was used, while a Tukey’s multiple comparisons test was used for **(B, D, and F)**. Asterisks indicate p-values of * = < 0.5, ** = < 0.02, *** = < 0.01, and **** = < 0.0001.

In addition to observing germination of the strains in response to two of the most efficient germinants, we also determined the response to a known competitive inhibitor of TA-mediated spore germination, CDCA. K_i_ values for each strain ranged from 0.10 mM for *C. difficile* PUC_75 to 0.90 mM for *C. difficile* strains PUC_256 and ICC5 (Figure 5E). The *C. difficile* R20291 control strain had a K_i_ value of 0.40 mM. There were no major differences in inhibitor sensitivity compared to the control. Additionally, this data is consistent between clades with only a 0.10 mM difference between the least (0.40 mM, Clade 3) and greatest (0.50 mM, Clade 1) K_i_ values (Figure 5F).

The bile acid and amino acid co-germinant signals are recognized in *C. difficile* by the pseudoproteases CspC and CspA respectively (46, 76). These signals are transmitted to CspB which then activates the cortex lytic enzyme, SleC (77–79). To determine if there was any genetic variation in the proteins responsible for initiating germination, we aligned the CspBA, CspC, and SleC sequences (46, 76, 77, 79, 80).

The alignments (Figure S10 – 12) of each protein revealed some variations compared to the *C. difficile* R20291 control strain, but none matched any residues which were found to influence germinant sensitivity (77, 80).

### *C. difficile* strains are equally resistant to bile acid toxicity

Several bile acids are known to inhibit *C. difficile* growth (75, 81, 82). Thus, we sought to determine if any of our strains were equally susceptible to CA, DCA, and CDCA using MIC assays. To mimic the various regions of the colon encountered by *C. difficile*, each assay was performed at pH 7.5, 6.8, 6.2, and 5.5 (83–85). The results indicated little variation in bile acid sensitivity between strains (Figure 6). At the most neutral pH, the MIC for CA was 7.5 mM. As the pH became more acidic, the MIC decreased 8-fold to 0.94 mM (Figure 6A). A similar trend was seen for CDCA and DCA with an MIC of 1 mM at pH 7.5, and an 8-fold decrease in MIC at pH 5.5 (Figure 6C, E). This data, when grouped by clade again found no difference in bile acid sensitivity between clades (Figure 6B, D, F).

**Figure 6:**
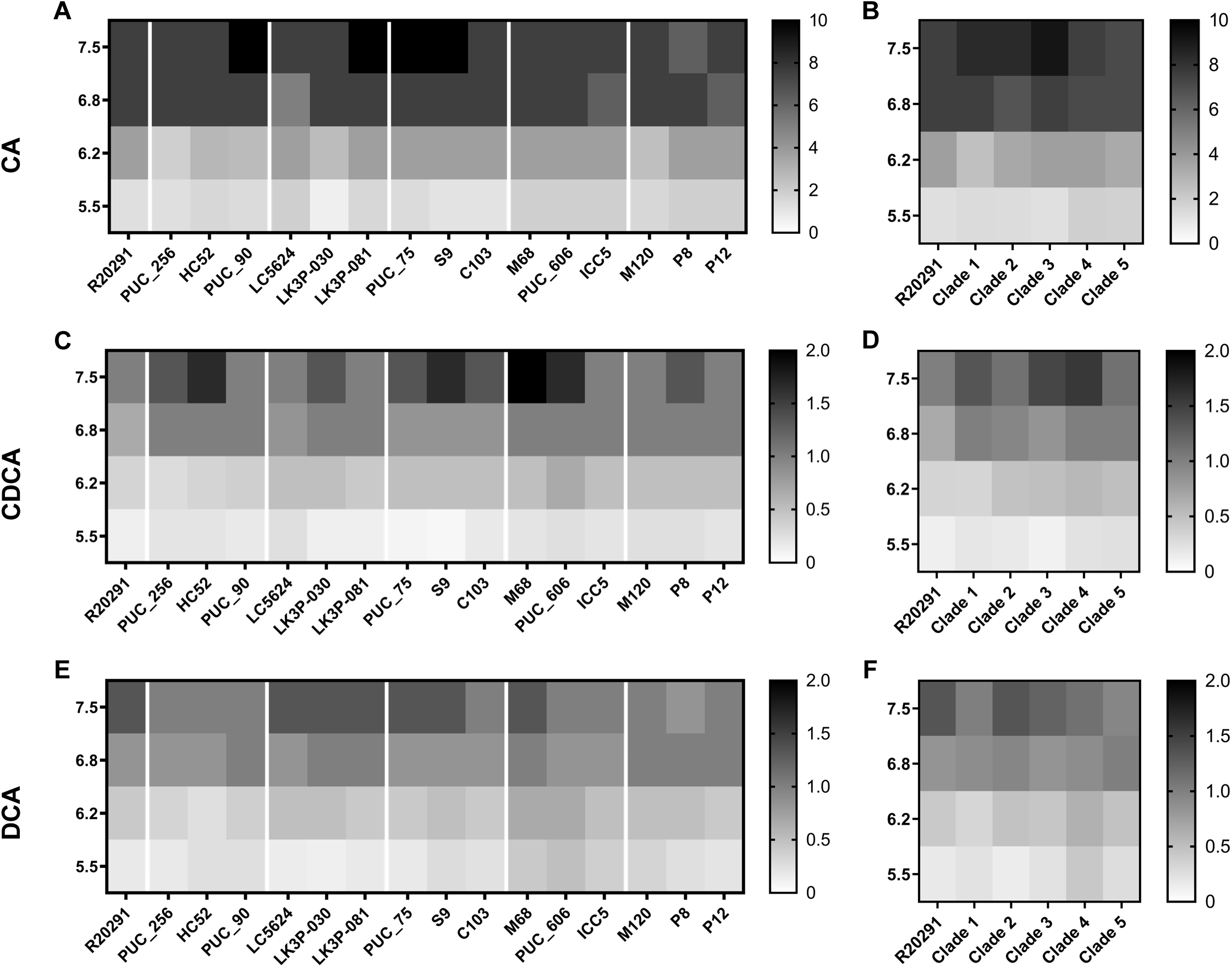
Bile salt sensitivity of strains. Stationary-phase cultures of each strain were inoculated into BHIS of the indicated pH and concentration of CA **(A,B)**, CDCA **(C,D)**, and DCA **(E,F)**. MICs were assessed by the presence/absence of growth after ∼18 hours. Data represents the average from independent biological triplicates

### *C. difficile* strains vary in their ability to modify taurine-based bile salts

Previous work from our lab demonstrated that some *C. difficile* strains deconjugate taurine-conjugated bile salts (48). While it is currently unknown how this activity contributes to disease outcomes, introduction of free taurine into the host could impact disease progression. Because the examined strains had various levels of activity, we tested the bile salt hydrolase activity (BSHA) of our strains using TA as a substrate. Each strain was incubated with TA and the amount of its deconjugated product (CA) was quantified (Figure 7). *C. difficile* PUC_90 (Clade 1) and *C. difficile* ICC5 (Clade 4) processed TA at an efficiency comparable to the *C. difficile* control (∼70 – 80%). *C. difficile* strains HC52 (Clade 1), S9 (Clade 3), and PUC_606 (Clade 4) had little activity against TA, while the remaining Clade 1, 3, and 4 strains had some activity (though not to the same level as *C. difficile* R20291). Interestingly, all non-control strains from both Clades 2 and 5 had low levels of BSHA against TA. This is surprising given that *C. difficile* R20291 also belongs to Clade 2.

**Figure 7:**
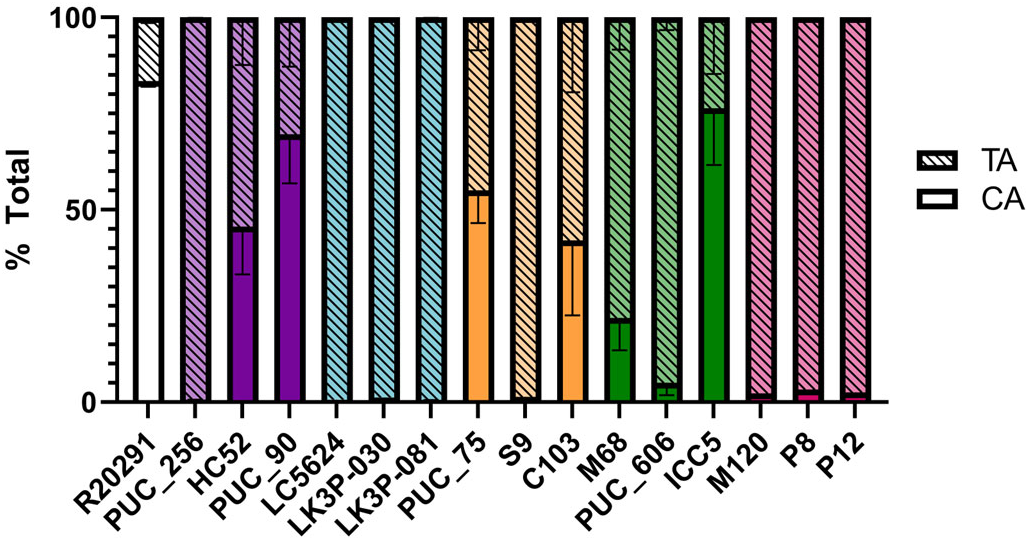
Bile salt hydrolase activity. Each strain was grown in the presence of 1 mM TA and incubated for 24 hours. The bile salts present in each culture following incubation were identified / quantified by reverse-phase high performance liquid chromatography (HPLC). Percent deconjugation was calculated using the following formula: % deconjugation = CA / (TA+CA). Data points represent the average from independent biological triplicates with error bars representing the standard error of the mean.

In prior work we found that strains which did not deconjugate TA could process TDCA (48). Thus, we tested if the Clade 2 and 5 strains could deconjugate TDCA (Figure S13). No detectable amount of DCA was generated by these strains. Taken together, this suggests that BSH activity may not be a core phenotype shared among strains.

### *C. difficile* strains show similar surface motility

Some Clade 5 strains have distinct growth morphologies compared to members of other clades (60). We tested surface motility for each of the strains to determine if we could detect similar morphologies (Figure S14). Strains from all clades (including the *C. difficile* R20291 control) spread similarly from the point of inoculation. The size of the protrusions from the central growth ring were the most striking variations, with *C. difficile* LK3P-030 (Clade 2) having the largest projections, and *C. difficile* M68 (Clade 4) producing virtually no projections. As previously reported (60), the clade 5 strains had some asymmetric growth of the extensions which is especially evident in the *C. difficile* P12 strain (Figure S14Fii).

## Discussion

### *C. difficile* is a highly variable species with an open pan-genome (18)

Phylogenetic analysis indicates significant evolutionary changes between the five main clades as indicated by widespread homologous recombination and horizontal gene transfer (19, 20). Previous research has shown that even members of the same clade / ribotype had varying phenotypes (*e.g.*, toxin activity) (86, 87). Additionally, several studies have observed an apparent correlation between ribotype and clinical outcome with RT106 (Clade 2) and RT078 (Clade 5) being associated with the worst outcomes / recurrence compared to the commonly studied RT027 (Clade 2) (30–38). We hypothesized that despite evolutionary variation, *C. difficile* strains would have core phenotypes that are central to *C. difficile* biology and consequently similar across clades / ribotyptes.

All strains had similar growth rates in the rich medium tested, indicating that any differences observed between strains is not due to differences in growth. When growth was assessed in a minimal medium, and compared to the laboratory *C. difficile* R20291 strain, the clinical strains showed decreased growth rates in glucose containing CDMM and increased growth rates in fructose / xylose / trehalose containing medium. Between the strains however, there was little variation, except for the Clade 4 strains which had consistently decreased growth rates in all tested media types. While alignments of the carbohydrate metabolism protein sequences revealed some genetic variation, these variations are not consistent with the observed phenotypic differences. This suggests that variations within the protein sequences themselves are not sufficient to explain the observed phenotypic variation.

Notably, another study reported that RT023 (Clade 3 strains) were unable to grow on CDMM-tre (88). Our Clade 3 strains grew relatively well in CDMM-tre which may be the result of differences in experimental set up. Specifically, the strains in this study were grown in CDMM-tre for 24 hours followed by a subculture into the same media and a further 12 hours of growth prior to inoculation of the experimental culture. In the Midani study (88), cells were grown solely in BHIS prior to dilution into the experimental culture. If trehalose metabolism occurs in these strains via an unconventional pathway as suggested by our genomic data, processing of trehalose may be less efficient during the lag phase of growth. The additional time spent in the trehalose-supplemented minimal medium likely allowed the strains to adapt to the nutrient limited conditions, reducing the lag-phase and resulting in growth observable over 22 hours.

For the number of spores produced by each strain over a 48-hour period, only two strains (*C. difficile* HC52 – Clade 1 and *C. difficile* LK3P-081 – Clade 2) demonstrated any statistically significant differences compared to *C. difficile* R20291; the largest difference observed between the strains was ∼10-fold. Whether this difference has any biological relevance remains to be seen. Given that some studies of *C. difficile* infection in mice report disease development with as few as 100 spores, this difference is unlikely to impact disease formation / progression but might contribute to altered disease spread / recurrence (6, 89).

We observed a decreased EC_50_ value for both TA and glycine for most of our strains compared to the control. This indicates increased sensitivity to these germinants. The EC_50_ of each germinant is less than or similar to its physiologically relevant concentration, suggesting that these strains have become better adapted to the mammalian gut (90, 91). Additionally, we observed no differences in germination in the presence of CDCA, a competitive inhibitor of TA mediated spore germination. CDCA was an effective inhibitor of TA-mediated germination in all tested strains. These results suggest anti-germination-based therapies could be broadly applicable in the treatment of CDI. In support of this, recent work on the bile salt analogs, CamSA & CaPA, show protection against CDI in mouse models of infection (92, 93).

Each of the strains was assessed for its ability to resist the toxic effects of certain bile acids. This assay was performed at various pHs to mimic conditions experienced by vegetative cells in the colon (83–85). We observed little variation in MIC when compared to *C. difficile* R20291 or between strains. As expected, the MIC of each bile salt increased with pH, reflecting the deprotonation of each bile acid (more negative charges) which limits its ability to interact with and disrupt the negatively charged bacterial membrane. The lack of variation here suggests that bile acid toxicity is a core phenotype for *C. difficile*. Moreover, our results suggest that these bile acids may have different effects on *C. difficile* growth depending on the location within the gut.

Recent work from our lab found that *C. difficile* is capable of processing taurine-conjugated bile salts by removing the taurine group (48). When this assay was performed on the strains in this study, they showed varying abilities to deconjugate TA, with Clade 2 and 5 members showing low processing levels. These same strains could also not process TDCA, indicating that the proteins responsible for this behavior are either missing, are not expressed, or have different substrates than the two tested herein. Interestingly, this phenotype is unique to the Clade 2 and 5 strains (not including the *C. difficile* R20291 control) which we noted previously have been associated with more severe / recurrent CDI. Because a bile salt hydrolase has not yet been identified in *C. difficile*, it remains unclear if BSHA, or the lack thereof, is relevant to the clinical outcome of CDI. Regardless, because not all the tested strains demonstrated BSHA, this may not be a core phenotype.

Most of the variation seen within each assay is observed in comparison to the control strain, *C. difficile* R20291. This strain was isolated during the Stoke-Manderville outbreak in the early 2000’s, has been passed between laboratories, and, likely, has since become a laboratory-adapted strain (though still virulent in animal models) (15). Observed differences correspond to an increased fitness of the clinical isolates compared to the *C. difficile* R20291 strain as indicated by increased growth rates in fructose / xylose / trehalose containing medium and increased sensitivity to germinants. This could indicate either a loss of some functions within the *C. difficile* R20291 strain or that *C. difficile*, as a species, has evolved to become more fit in the gut.

When considering the strains independently of *C. difficile* R20291, we observed remarkable phenotypic similarity between strains and no major patterns corresponding to clade / ribotype. This is true of every tested phenotype except for BSHA, suggesting that it may not be a core process; only identifying the factor responsible for BSHA will test this hypothesis. Taken together, this data suggests that the previously observed relationships between ribotype and CDI severity may not be due to changes in these core phenotypes but rather to other influences.

This study focused only on a small portion of the known *C. difficile* strains, and a limited number of phenotypes. Further analysis of phenotypic variation between strains of all ribotypes / clades both *in vivo* and *in vitro* will expand upon what we have learned here and provide valuable insight into how *C. difficile* might manifest itself in a clinical setting.

## Acknowledgments

Thanks to the members of the Sorg laboratory for their helpful comments during the preparation of this manuscript. This project was supported by awards 5R01AI116895 and 5R01AI172043 to J.A.S. and 1R01AI177842 to D.P.S from the National Institute of Allergy and Infectious Diseases. The content is solely the responsibility of the authors and does not necessarily represent the official views of the NIAID. The funders had no role in study design, data collection and interpretation, or the decision to submit the work for publication.

## Supplemental Figures

**Figure S1: *C. difficile* R20291 controls and EC_50,glycine_ assay**

A) Germination of purified R20291 spores under control conditions; buffer only, TA only, Gly + DMSO, and TA + Gly + DMSO. These controls were run on every germination assay plate. B) Germination of purified R20291 spores exposed to the indicated concentrations of glycine.

**Figure S2: *C. difficile* R20291 BSHA controls**

Chromatograms for R20291 without bile salt treatment (A), with TA, HDCA, and CA added after centrifugation (B), and with TA added prior to the 24 hour incubation (C). The chromatograms from A-C are overlayed in (D) to confirm the identity of the peaks.

**Figure S3: Additional phylogenies**

The neighbor joining phylogeny generated for the strains in this study created from LCBs 117, a 842,010 bp segment (A) and 71, a 228,563 bp segment extracted from the MAUVE alignment (B). The phylogeny was constructed using the Geneious Tree Builder application in Geneious Prime software using the Tamura-Nei genetic distance model. Strains are grouped by their respective ribotypes / clades with the scale bar representing the number of substitutions per 1000 bp.

**Figure S4: ClustalOmega alignment of CcpA**

The gene sequence of *ccpA* was extracted from the MAUVE alignment. The sequences were translated and aligned using ClustalOmega. Similarity to the *C. difficile* R20291 control is indicated by shading with white indicating 100% similarity and black indicating <60 % identity.

**Figure S5: ClustalOmega alignment of XylA**

The gene sequence of *xylA* was extracted from the MAUVE alignment. The sequences were translated and aligned using ClustalOmega. Similarity to the *C. difficile* R20291 control is indicated by shading with white indicating 100% similarity and black indicating <60 % identity.

**Figure S6: ClustalOmega alignment of XylB**

The gene sequence of *xylB* was extracted from the MAUVE alignment. The sequences were translated and aligned using ClustalOmega. Similarity to the *C. difficile* R20291 control is indicated by shading with white indicating 100% similarity and black indicating <60 % identity.

**Figure S7: ClustalOmega alignment of XylR**

The gene sequence of *xylR* was extracted from the MAUVE alignment. The sequences were translated and aligned using ClustalOmega. Similarity to the *C. difficile* R20291 control is indicated by shading with white indicating 100% similarity and black indicating <60 % identity.

**Figure S8: ClustalOmega alignment of TreA**

The gene sequence of *treA* was extracted from the MAUVE alignment. The sequences were translated and aligned using ClustalOmega. Similarity to the *C. difficile* R20291 control is indicated by shading with white indicating 100% similarity and black indicating <60 % identity.

**Figure S9: ClustalOmega alignment of TreR**

The gene sequence of *treR* was extracted from the MAUVE alignment. The sequences were translated and aligned using ClustalOmega. Similarity to the *C. difficile* R20291 control is indicated by shading with white indicating 100% similarity and black indicating <60 % identity.

**Figure S10: ClustalOmega alignment of CspBA**

The gene sequence of *cspBA* was extracted from the MAUVE alignment. The sequences were translated and aligned using ClustalOmega. Similarity to the *C. difficile* R20291 control is indicated by shading with white indicating 100% similarity and black indicating <60 % identity.

**Figure S11: ClustalOmega alignment of CspC**

The gene sequence of *cspC* was extracted from the MAUVE alignment. The sequences were translated and aligned using ClustalOmega. Similarity to the *C. difficile* R20291 control is indicated by shading with white indicating 100% similarity and black indicating <60 % identity.

**Figure S12: ClustalOmega alignment of SleC**

The gene sequence of *sleC* was extracted from the MAUVE alignment. The sequences were translated and aligned using ClustalOmega. Similarity to the *C. difficile* R20291 control is indicated by shading with white indicating 100% similarity and black indicating <60 % identity.

**Figure S13 Bile salt hydrolase activity - TDCA**

Each strain was grown in the presence of 1 mM TDCA and incubated for 24 hours. The bile salts present in each culture following incubation were identified / quantified by reverse-phase high performance liquid chromatography (HPLC). Percent deconjugation was calculated using the following formula: % deconjugation = DCA / (TDCA+DCA). Data points represent the average from independent biological triplicates with error bars representing the standard error of the mean.

**Figure S14: Surface motility assay**

Stationary phase cultures were spotted into BHIS and incubated for 5 days prior to imaging. Each panel is a representative image from three distinct biological replicates.

A) *C. difficile* R20291. B) Clade 1-RT024-020 strains: Bi) *C. difficile* PUC_256, Bii) *C. difficile* HC52, and Biii) *C. difficile* PUC_90. C) Clade 2-RT106 strains: Ci) *C. difficile* LC5624, Cii) *C. difficile* LK3P-030, and Ciii) *C. difficile*. LK3P-081. D) Clade 3-RT023 strains: Di) *C. difficile* PUC_75, Dii) *C. difficile* S9, and Diii) *C. difficile* C103. E) Clade 4- RT017 strains: Ei) *C. difficile* M68, Eii) *C. difficile* PUC_606, and Eiii) *C. difficile* ICC5. F) Clade 5-RT078 strains: Fi) *C. difficile* M120, Fii) *C. difficile* P8, and Fiii) *C. difficile* P12.

